# A HUMANIZED FCRN TRANSGENIC MOUSE FOR PRECLINICAL PHARMACOKINETICS STUDIES

**DOI:** 10.1101/2022.10.24.513622

**Authors:** Christopher M. Conner, Don van Fossan, Kristen Read, Dale O. Cowley, Oscar Alvarez, Shannon (Xiang-Ru) Xu, David R. Webb, Kurt Jarnagin

## Abstract

Monoclonal antibodies (mAbs) are one of the fastest-growing classes of drugs and have been approved to treat several diseases, including cancers and autoimmune disorders. Preclinical pharmacokinetics studies are performed to determine the therapeutically meaningful dosages and efficacy of candidate drugs. These studies are typically performed in non-human primates; however, using primates is costly and raises ethical considerations. As a result, rodent models that better mimic human-like pharmacokinetics have been generated and remain an area of active investigation.

Pharmacokinetic characteristics of a candidate drug, such as half-life, are partly controlled by antibody binding to the human neonatal receptor hFCRN. Due to the abnormally high binding of human antibodies to mouse FCRN, traditional laboratory rodents do not accurately model the pharmacokinetics of human mAbs. In response, humanized rodents expressing h*FCRN* have been generated. However, these models generally use large inserts randomly integrated into the mouse genome. Here, we report the production and characterization of a CRISPR/Cas9-mediated h*FCRN* transgenic mouse termed SYNB-h*FCRN*. Using CRISPR/Cas9-assisted gene targeting, we prepared a strain with a simultaneous knockout of m*Fcrn* and insertion of a h*FCRN* mini-gene under the control of the endogenous mouse promoter. These mice are healthy and express hFCRN in the appropriate tissues and immune cell subtypes. Pharmacokinetic evaluation of human IgG and adalimumab (Humira®) demonstrate h*FCRN-* mediated protection. These newly generated SYNB-h*FCRN* mice provide another valuable animal model for use in preclinical pharmacokinetics studies during early drug development.

**Materials availability:** Cryopreserved germline cells from the strains are available from the MMRRC. Dr. Jarnagin will provide sequencing and other evidence supporting strain identity and the plasmids used for constructions.

## Introduction

Therapeutic monoclonal antibodies (mAbs) are a growing class of drugs with more than one hundred approved by the United States Food and Drug Administration (FDA). These mAbs are approved to treat various diseases, including immunological disorders and cancers (Kaplon et al., 2022). During drug development, preclinical pharmacokinetics (PK) studies are often performed using non-human primates (NHP). However, NHP are less accessible, expensive and raise ethical concerns. A rodent model of human PK would increase the animal model availability, reduce the cost, and moderate ethical concerns, thus improving mAb drug development.

Understanding the PK of newly developed mAbs is critical for achieving therapeutically meaningful exposure and therapeutic efficacy. Preclinical research is often aimed at lengthening antibody half-lives to improve exposure and efficacy; hence inexpensive, readily available models are needed. Most therapeutic mAbs are immunoglobulin G (IgG) subclasses, IgG1 and IgG2 (Ryman and Meibohm, 2017). The neonatal Fc receptor (FCRN; also known as FCGRT) plays a central role in IgG clearance by protecting antibodies from degradation, thus prolonging their PK (Ahouse et al., 1993; Chaudhury et al., 2003; Simister et al., 1989; Roopenian et al., 2003). However, there are species differences in antibody binding to FCRN (Ober et al., 2001). For example, human IgG has a higher binding affinity for mouse FCRN (mFCRN) than for human FCRN (hFCRN) (Ober et al., 2001). This abnormally high affinity for mFCRN reduces the utility of wildtype (WT) mice in therapeutic mAb development (Proetzel et al., 2013).

To address the cross-species differences, laboratories have generated mice lacking the *Fcrn gene* and expressing h*FCRN*. Two widely used “humanized” mouse strains are the Tg32 strain using the human promoter to drive the expression of a randomly integrated transgene, and the Tg276 strain using the strong-promiscuous chicken β-actin promoter to drive expression (Roopenian et al., 2010). Consistent with the promoters employed, the tissue expression pattern of hFCRN in Tg32 mice is similar to humans and monkeys, whereas Tg276 mice express higher levels of hFCRN in many tissues (Latvala et al., 2017). These findings have led to the understanding that Tg32 mice are better suited for preclinical PK studies than Tg276. Several reports demonstrate its utility in predicting human PK (Tam et al., 2013; Avery et al., 2016; Valente et al., 2020). However, the Tg32 strain was generated using a large insert (34kb) with random integration at chromosome 2 (Roopenian et al., 2010). Additionally, the transgene is under the control of the native human promoter, which may not function optimally in the mouse cellular environment. We created a new hFCRN transgenic mouse to address these issues using CRISPR/Cas9-assisted gene targeting (Ittner and Gotz, 2007; Li et al., 2013; Wang et al., 2013; Lai et al., 2022). The *SYNB-hFCRN* line has a h*FCRN* mini-gene (4kb) inserted into the m*Fcrn* locus on mouse chromosome 7 such that the hFCRN is under the control of the mouse promoter while simultaneously producing a knockout (KO) of the m*Fcrn* gene.

Here, we report the production and characterization of SYNB-h*FCRN* by first confirming RNA and protein expression at the tissue and cellular levels. We next measured levels of endogenous mouse IgG to establish the effect of hFCRN on this plasma protein. After characterization of the knockout and expression patterns of the human mRNA and protein of SYNB-h*FCRN*, we functionally evaluated hFCRN by comparing the PK of human IgG between wildtype (WT), m*Fcrn* KO, Tg32 h*FCRN*, and SYNB-h*FCRN* mice. We also evaluated the PK of one of the most widely used therapeutic mAbs, adalimumab (Humira®).

## Results

### Generation and health of SYNB-h*FCRN* mice

To generate a m*Fcrn* deficient, h*FCRN* knock-in (KI) animal, we used CRISPR/Cas9-assisted gene targeting to create a KO of m*Fcrn* while simultaneously inserting a h*FCRN* mini-gene under the control of the mouse promoter (Fig. 1A). The SYNB-h*FCRN* mice appear normal and showed no clinically relevant or statistically significant differences in body weight or spleen weight compared to wildtype (WT) mice (Fig. 1B). We confirmed KO of m*Fcrn* and KI of h*FCRN* by RT-PCR analysis of mRNA from the SYNB-h*FCRN* strain (Fig. 1C). In organs associated with FCRN function, SYNB-h*FCRN* mice express h*FCRN* in the spleen, liver, kidney, and intestine matching the tissue expression pattern observed for m*Fcrn* in WT tissues (Fig. 1C). These data demonstrated that KO of m*Fcrn* and insertion of h*FCRN* was successful. RNA expression of h*FCRN* occurred in the expected tissues. We performed flow cytometric analysis on splenocytes harvested from SYNB-h*FCRN* mice (Fig. 2A). We observed hFCRN expression in monocytes, natural killer (NK) cells, and macrophages (Fig. 2B and 2C). The data demonstrate that expression of hFCRN occurs in the proper immune cell subpopulations.

**Figure 1.**
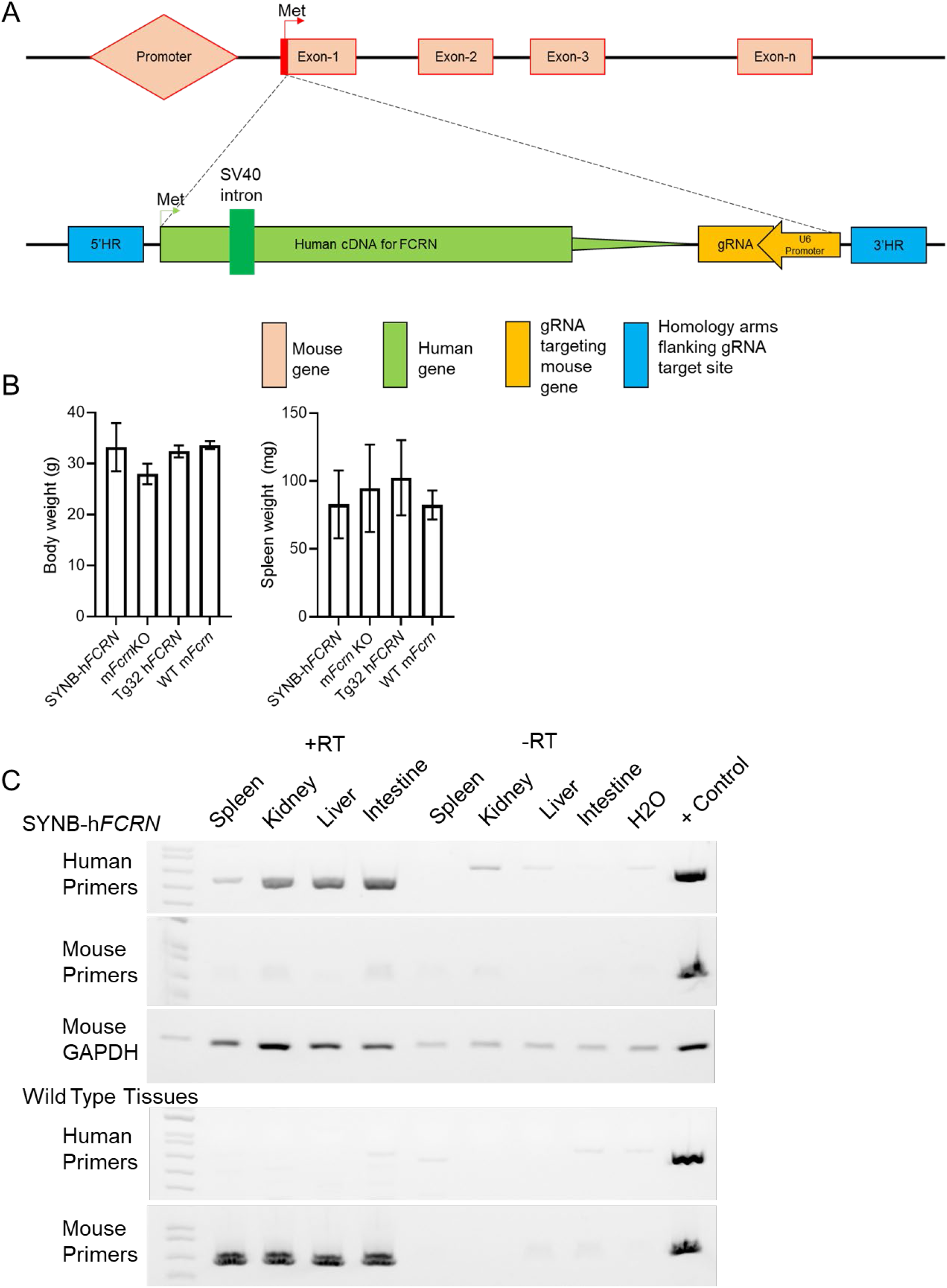
Generation and health of SYNB-h*FCRN* mice. (A) Gene insertion schematic of cDNA for h*FCRN* at the mouse *Fcgr* locus. (B) Body weights and spleen weights of 13w old SYNB-h*FCRN*, m*Fcrn* KO, Tg32 h*FCRN*, and wildtype mice. Error bars denote SD, 3 – 4 individual mice were weighed. (C). Tissue level gene expression of h*FCRN* mRNA in SYNB-h*FCRN* mice and wildtype mice.

**Figure 2.**
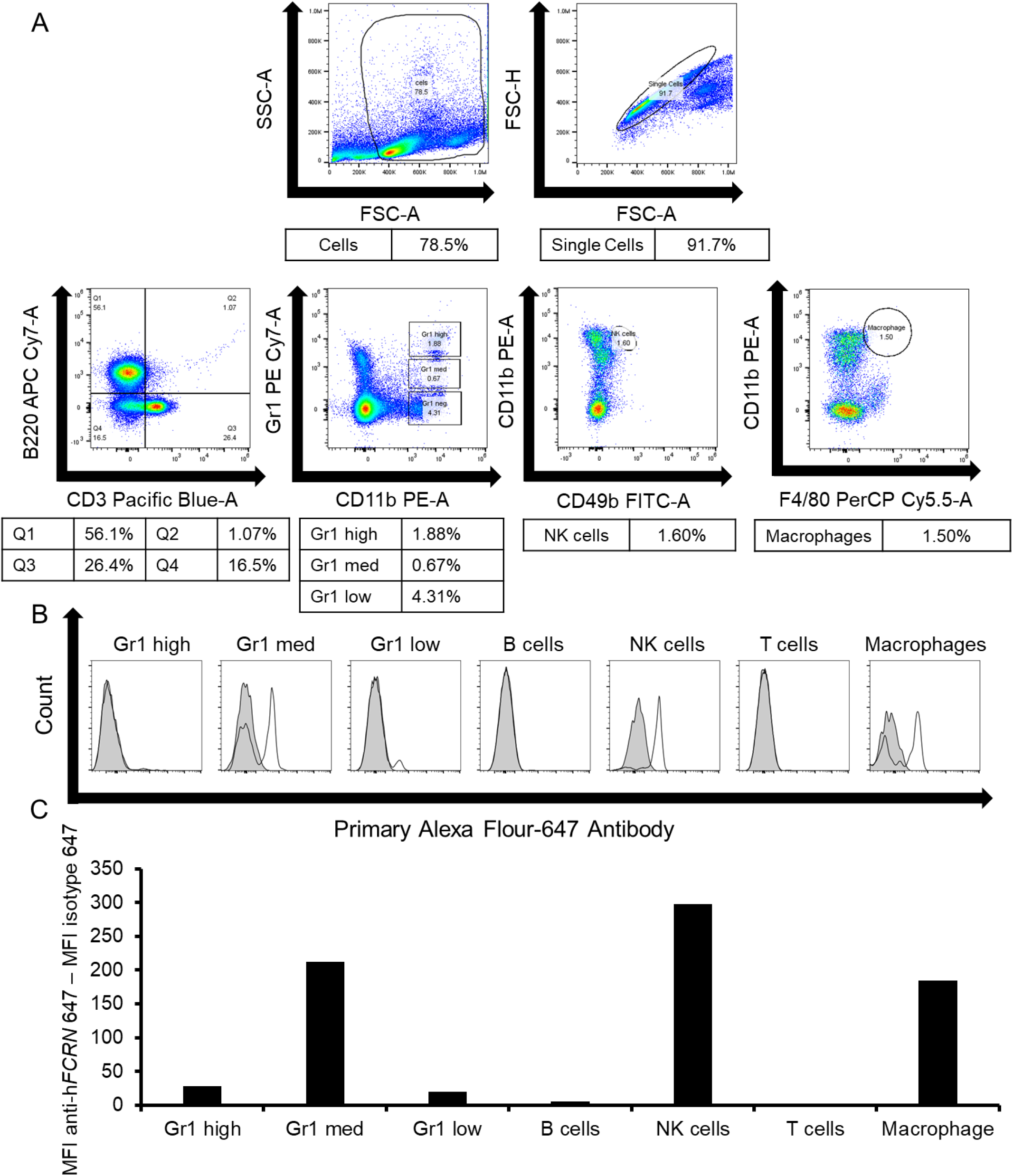
Flow cytometric analysis of hFCRN protein expression in SYNB-h*FCRN* mice. (A) Flow cytometric gating scheme to identify immune cell subtypes. (B). Histograms showing signal from anti-hFCRN-647 antibody (white histograms) overlayed with signal from isotype control antibody-647 (grey histograms). (C) Mean fluorescence intensity (MFI) of hFCRN signal subtract MFI from isotype control antibody. Gr1 high: neutrophils. Gr1 med to Gr1 low: monocytic. Data derived from a representative animal.

### Endogenous mouse IgG levels in SYNB-h*FCRN* mice

Endogenous mouse IgG has a lower binding affinity for hFCRN when compared to human IgG, which results in lower mIgG levels in Tg32 h*FCRN* mice compared to WT mice (Ober et al., 2001; Tam et al., 2013). To determine the effect of h*FCRN* expression on mouse IgG levels in SYNB-h*FCRN* mice, we measured IgG levels in WT, Tg32 h*FCRN*, m*Fcrn* KO, and SYNB-h*FCRN* mice. We observed lower levels of mouse IgG, by ~70%, in SYNB-h*FCRN* mice compared to WT mice (Fig. 3). This result indicates the successful removal of m*Fcrn* and the expected effects of h*FCRN* expression on endogenous mouse IgG levels.

**Figure 3.**
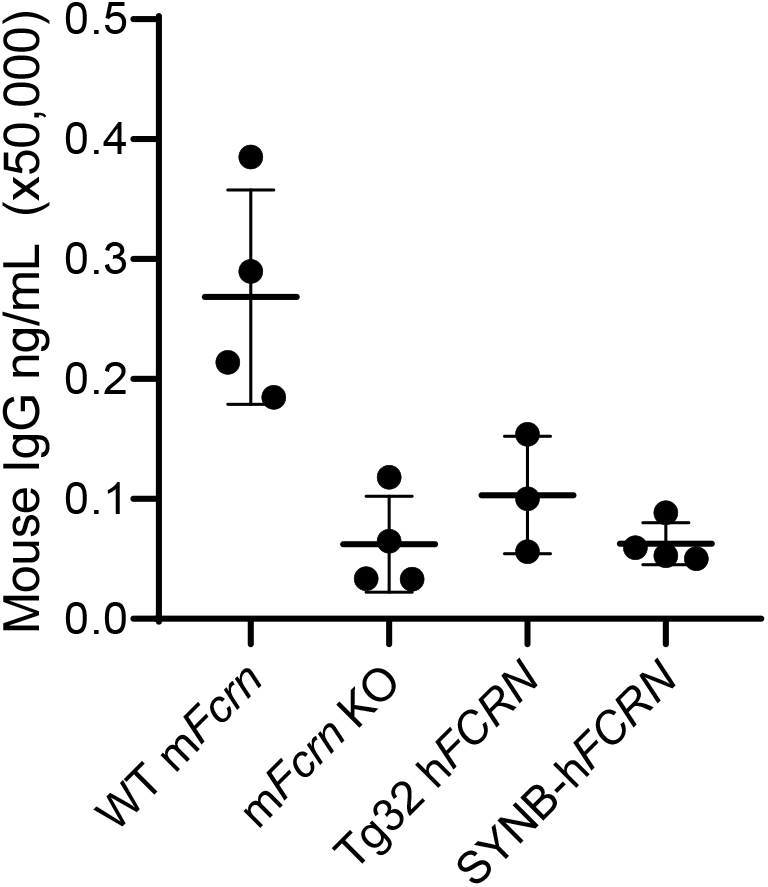
Mouse IgG levels. Mouse IgG levels in WT, m*Fcrn* KO, Tg32 h*FCRN*, and SYNB-h*FCRN* mice. Each dot represents an individual mouse. Horizontal line denotes mean and error bars denote SD. 3 - 4 mice were used per genotype.

### Human IgG pharmacokinetics in SYNB-h*FCRN* mice

Human IgG interaction with FCRN is critical for prolonging IgG half-life. To functionally characterize SYNB-h*FCRN* mice and compare other strains, we determined whether hFCRN expression protects human IgG from degradation. To test this, we injected cohorts of WT, Tg32 h*FCRN*, m*Fcrn* KO, and SYNB-h*FCRN* mice with pooled human IgG at 10 μg/g body weight. Mice were then serially bled via submandibular vein puncture starting 3 hours post-injection through 21 days post-injection. We observed no significant differences in human IgG PK between Tg32 h*FCRN* and SYNB-h*FCRN* mice (Fig. 4A). Additionally, we did not observe significant differences between WT and SYNB-h*FCRN* mice (Fig. 4B). In WT mice, the t_½_ was 15.55 days. The t_½_ in SYNB-h*FCRN* and Tg32 h*FCRN* mice was 11.98 and 13.52 days, respectively. As expected, human IgG degrades rapidly in m*Fcrn* KO mice resulting in a t_½_ of 0.97 days. These data indicate that hFCRN in SYNB-h*FCRN* mice is functional and protects human IgG from degradation.

**Figure 4.**
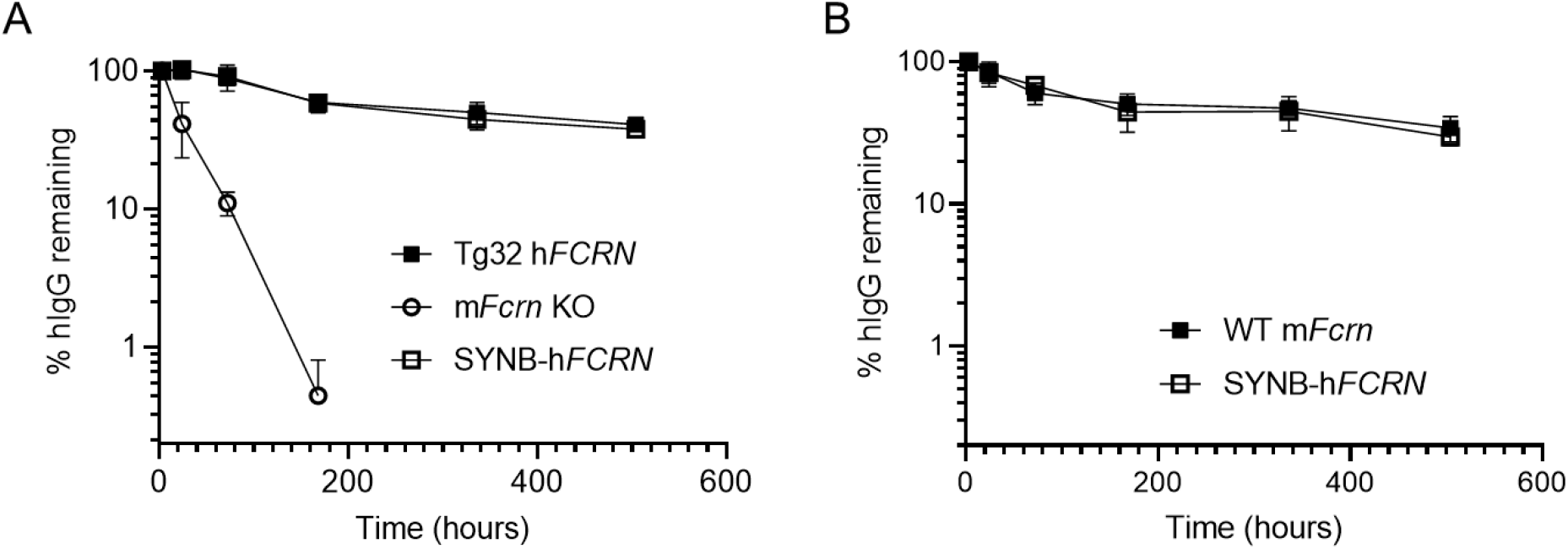
Human IgG pharmacokinetics. (A) PK profile of pooled human IgG in Tg32 h*FCRN*, m*Fcrn* KO, and SYNB-h*FCRN* mice. (B) PK profile of human IgG in WT and SYNB-h*FCRN* mice. Each dot represents 3 – 5 individual mice. Points are normalized to first bleed at 3 hours. Error bars denote SD.

### Evaluation of adalimumab (Humira®) pharmacokinetics in SYNB-h*FCRN* mice

To evaluate if hFCRN in SYNB-h*FCRN* mice protects clinically used mAbs, we chose to test the anti-TNFα biologic adalimumab (Humira^®^). Humira^®^ represents one of the most prescribed biologics in the United States. Adalimumab was administered using a single intraperitoneal dose at 2.5 mg/kg body weight. PK profiles were determined for 21 days in WT, Tg32 h*FCRN, mFcrn* KO, and SYNB-h*FCRN* mice (Fig. 5). Consistent with Myzithras et al., 2016; we observed rapid clearance of adalimumab within seven days (Fig. 5). Adalimumab exhibited an approximate t_½_ of 5.2 days in WT mice consistent with previously published work (Myzithras et al., 2016). PK of adalimumab was similar between SYNB-h*FCRN* and Tg32 h*FCRN* mice showing an approximate t_½_ of 2.08 days. Adalimumab is cleared faster in mFcrn KO mice with an approximate t_½_ of 0.833 days (Fig. 5). This data demonstrates that SYNB-hFCRN mice prolong adalimumab presence in the blood compared to knockout strains and provides human-like antibody-Fc receptor interactions and clearance.

**Figure 5.**
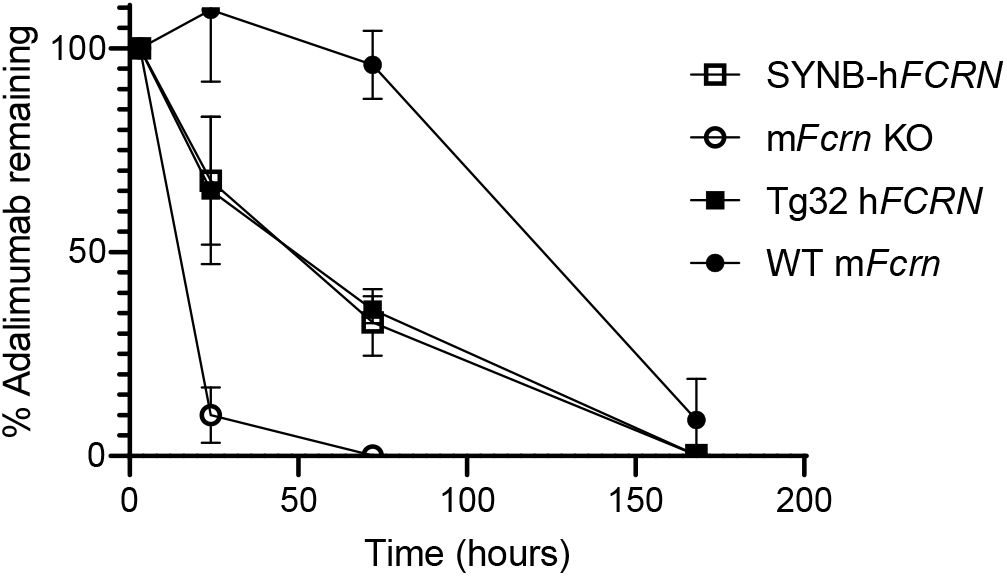
Adalimumab pharmacokinetics. PK profile of adalimumab in WT m*Fcrn*, Tg32 h*FCRN*, m*Fcrn* KO, and SYNB-h*FCRN* mice. Each dot represents 3 – 4 individual mice. Points are normalized to first bleed at 3 hours. Error bars denote SD.

## Discussion

Monoclonal antibodies and other biologics represent one of the fastest-growing drug classes in medicine. The development of therapeutic monoclonal antibodies requires preclinical pharmacokinetic studies. The NHP remains the gold standard for preclinical pharmacokinetics; however, their expense, availability, and ethical considerations motivate the development of alternative models. A rodent model that better recapitulates human PK may address these concerns and improve their usefulness in drug discovery efforts.

Most clinically used mAbs are IgG subclasses, primarily IgG1 and IgG2 (Ryman and Meibohm, 2017). These subtypes are selected partly due to the FCRN-mediated prolongation of the serum half-life of these IgG subtypes. As a result, several labs have generated “humanized mice” expressing hFCRN to serve as preclinical pharmacokinetics research models. However, the rodent models to date were developed using random integration of large inserts and may be under the control of promiscuous promoters driving artificially high levels of hFCRN expression. Herein, we used CRISPR/Cas-9 technology to generate a precisely humanized *FCRN* mouse called SYNB-h*FCRN*. Sequencing and RT-PCR data demonstrate successful knockout of mouse *Fcrn* and knock-in of human *FCRN* in the expected tissues. We report detection of hFCRN protein in the expected immune cell subpopulations, highest in the Gr+ medium monocytes, NK cells and macrophages. Removal of m*Fcrn* and expression of h*FCRN* resulted in reduced levels of endogenous mouse IgG levels, likely due to its weaker interaction with mouse IgG. FCRN in SYNB-h*FCRN* mice is functional and protects human IgG from degradation and prolongs the PK of adalimumab compared to m*Fcrn* KO animals. Our data comparing human IgG PK in WT and Tg32 h*FCRN* mice is consistent with previous studies (Roopenian et al., 2003; Roopenian et al., 2010).

Together these results demonstrate that SYNB-h*FCRN* mice are healthy and express functional hFCRN in the appropriate tissues. The h*FCRN* functions to lengthen antibody half-life in the expected manner as compared to the m*Fcrn* knockout mouse. These precisely integrated h*FCRN* mice (SYNB-h*FCRN*) will be a valuable resource for preclinical pharmacokinetics studies.

## Materials and Methods

### Animal husbandry and production of transgenic mice

All *in vivo* studies were conducted in compliance with an Institutional Animal Care and Use Committee (IACUC) protocol between Synbal Inc., and BTS, Inc, San Diego, CA. Humanization of the m*Fcrn* target loci followed a standard model (Ittner and Gotz, 2007; Li et al., 2013; Wang et al., 2013). The mouse target locus was reconfigured to express the human cDNA at the target site in exon 1 of the m*Fcrn*. We aimed to insert the human initiation methionine and cDNA at, or close to, and in-frame with, the mouse initiation methionine. This construct inserts an in-frame human mini-gene in exon 1 of m*Fcrn* and disrupts mouse gene expression. The humanization cassette contains a guide RNA expression cassette with a U6 promoter element driving expression of the same gDNA used to insert the human mini-gene into the mouse genome, resulting in targeting the same site on the wildtype mouse chromosome (more details may be found in Lai et al., 2022). The gRNA and Active Genetic cassette were incorporated to facilitate gene conversion and rapid breeding using the Super Mendelian principle (Lai et al., 2022). This project was active during the creation of SYNB-h*FCRN*. The Active Genetic cassette has no effect in strains not also expressing Cas9.

### mRNA isolation and RT PCR

Tissues (spleen, intestine, liver, and kidney) were harvested from WT or transgenic mice and immediately placed in RNA-later. mRNA was prepared using the RNAqueous-4PCR kit (Thermo Fisher), and cDNA synthesis was performed using the High-Capacity RNA-to-cDNA kit (Thermo Fisher) following the manufacturer’s protocols. Samples were analyzed using PCR applying primers on the 5’ side of the insert and on the 3’ side of the artificial intron (see Table 1). Agarose sizing gels were used to detect expression, and DNA sequencing of these bands after extraction was used to verify band identity. Studies were conducted with and without the addition of reverse transcriptase to assess genomic DNA contamination. GAPDH serves as a cDNA preparation quality control and gel loading control.

**Table 1.**
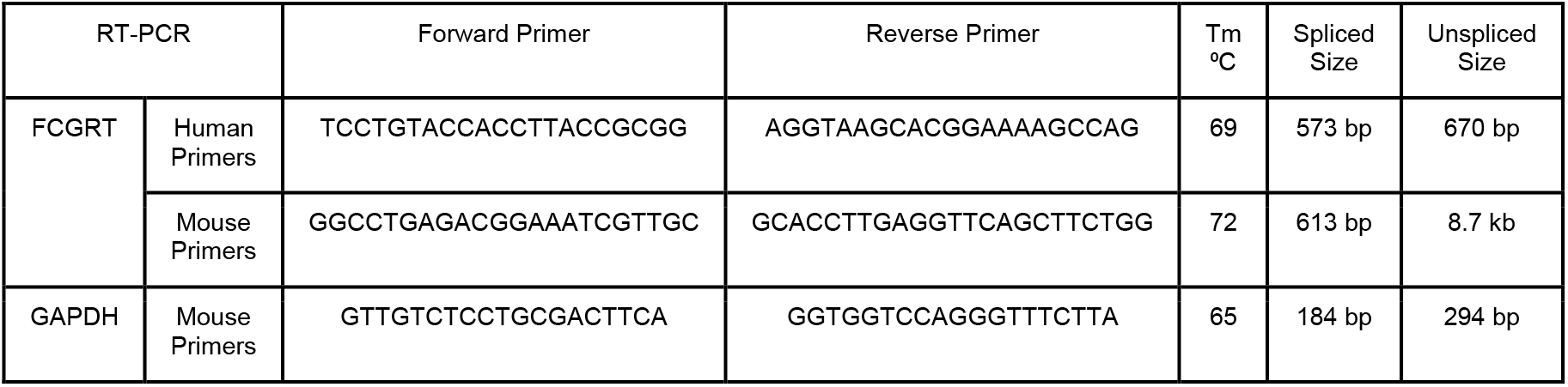
Primers

### Splenocyte preparation, antibody staining, and flow cytometry

Animals were sacrificed, and harvested spleens were placed on ice. Splenocytes were collected by pressing the spleens with the end of a 1mL syringe plunger through a 70μM filter (25-376, Genesee Scientific) in cold HBSS (14175079, Gibco). Filters were washed with 5mL cold HBSS to collect residual cells, and collection tubes were centrifuged. Cell pellets were resuspended in 1mL ammonium-chloride-potassium (ACK) red-cell lysis buffer (A1049201, Life Technologies) and kept on ice for 3 min. Cells were washed with 10mL cold HBSS, and pellets were resuspended in HBSS containing 2% heat-inactivated FBS (10082147, Gibco). All centrifugation steps were done at 1500 rpm for 5 min at 4°C.

Splenocyte cell suspensions (~2 million cells in HBSS) were incubated with Fc block (553142, BD Biosciences) for 10 min on ice followed by the direct addition of the following antibodies, Pacific Blue anti-mouse CD3 (100213, BioLegend), FITC anti-mouse CD49b (108905, BioLegend), APC/Cyanine7 anti-mouse/human CD45R/B220 (103223, BioLegend), PE/Cyanine7 anti-mouse Ly-6G/Ly-6C (108415, BioLegend), PE anti-mouse /human CD11b (101207, BioLegend), PerCP/Cyanine5.5 anti-mouse F4/80 (123127, BioLegend). Cells were incubated in antibodies for 30 min at 4°C in the dark. Samples were then washed twice with PBS and resuspended in 100μL Cytofix/Cytoperm (554722, BD Biosciences). Following fixation, cells were washed twice with intracellular staining permeabilization wash buffer (421002, BioLegend), followed by incubation with Alexa Fluor 647 labeled anti-human FCRN (IC8639R100U, RnD systems) for 30 min at 4°C in the dark. 400μL PBS was added to each sample, and flow cytometric analysis was performed on a Thermo Scientific Attune NxT Flow Cytometer. For compensation setup, single stain preparations of each antibody using splenocytes or UltraComp eBeads (01-3333-42, Life Technologies) were used. Alexa Flour 647 mouse IgG2b isotype control antibody (IC0041R, Novus Biologicals) and WT splenocytes were negative controls. All centrifugation steps were done at 2500 rpm for 3 min at room temperature. Data analysis and visualization were performed using FlowJo software (Version 10.7.2, BD). See table 2 for gating criteria.

**Table 2.** Flow cytometry gating criteria to identify immune cell subpopulations.

| Cell Population | Gating Criteria |
| --- | --- |
| Gr1 high(neutrophils) | Gr1+ ($>10^3$)/CD11b+ |
| Gr1 med (monocytic) | Gr1+ ($10^2 - 10^3$)/CD11b+ |
| Gr1 neg (monocytic) | Gr1-/CD11b+ |
| Macrophages | CD3-/B220-/CD11b+/F480+ |
| B cells | CD3-/B220+ |
| NK cells | CD3-/B220-/CD11b+/CD49b+ |
| T cells | CD3+/B220- |

### Pharmacokinetics studies

Pharmacokinetics studies were conducted in C57BL/6 mice (Jackson Labs Stock No. 000664), Tg32 (Jackson Labs Stock no. 014565), and mFcRn KO (Jackson Labs Stock No. 003982) mice, all purchased from The Jackson Laboratory (Bar Harbor, Maine). Transgenic mice expressing hFcRn were bred at Synbal Inc (San Diego, California). All mice were treatment naïve males between the ages of 8 to 20 weeks. For human IgG studies, mice were injected with 10 μg/g body weight in sterile PBS via intraperitoneal injection. In adalimumab (A2010, Selleck Chemicals) studies, mice were injected with 2.5mg/kg body weight in sterile PBS via intraperitoneal injection. Injection volumes were between 180μL and 210μL /mouse.

For each genotype, three subgroups of 4 to 5 age-matched mice were serially bled twice via submandibular vein puncture using 25-gauge needles. Mice were anesthetized with isoflurane and oxygen before bleeding. Mice were bled 3 h, 1 d, 3 d, 7 d, 14 d, and 21 d post-injection. Blood withdrawals were conducted so that each subgroup of mice was only bled once per week. For example, subgroup A was bled at the 3 hours and 1-week time points, subgroup B was bled at the 1-day and 2-week time points, and subgroup C was bled at the 3-day and 21-day time points. Blood samples (100μL - 200μL) were collected into EDTA-coated capillary collection tubes (Cat No., 20.1288.100, Sarstedt through ThermoFisher, Waltham, MA, USA) or tubes containing 2μL of 0.5M EDTA to prevent clotting. Blood samples were immediately placed on ice and spun down at 2,000 x g for 10 mins at room temperature, and the supernatant was collected into new 1.5mL tubes. Plasma was stored at −80°C until all samples were collected and analysis could be performed. Multiple freeze/thaw cycles were avoided. For mouse IgG studies, 3 to 4 age-matched mice were bled once via submandibular vein puncture.

### Enzyme-linked immunosorbent assay (ELISA)

Quantification of plasma levels of human IgG, mouse IgG, and adalimumab (Humira) was determined by an enzyme-linked immunosorbent assay (ELISA). For human and mouse IgG studies, plasma samples were thawed and diluted 1/400 and 1/50000, respectively, in ELISA buffer. ELISA was performed following the manufacturer’s protocols (Human IgG ELISA E88-104 and Mouse IgG ELISA E99-131, Bethyl Laboratories). For adalimumab studies, plasma samples were thawed and diluted 1/1000 in ELISA buffer, and ELISA was performed following the manufacturer’s protocol (Adalimumab ELISA KBI1015, Krishgen Biosystems). Following the addition of stop reagents, absorbance at 450nm was immediately read using a Promega GloMax Discover microplate reader (Madison, Wisconsin). Data analysis was performed using GraphPad Prism to fit standard curves into a 4-parameter logistic curve. Half-life was calculated using the following formula:

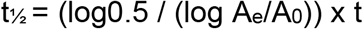

Where t_½_ is the molecule’s half-life, A_e_ is the amount of molecule remaining, A_0_ is the amount of molecule at the first bleed, and t is the elapsed time. When half-life could not be accurately calculated due to a lack of terminal time points, rapid elimination, or non-first order pharmacokinetics, the half-life was approximated by drawing a straight line from the 50% point on the Y axis (% molecular remaining) and interpolating a value on the X axis (time in hours).

## Acknowledgments

Synbal would like to thank Dr. Stephen Hedrick for providing valuable input. The interest and support of Drs. Ho Cho, Kate Blease, Rama Narla, and Winston Thomas are gratefully acknowledged. TransViragen would like to acknowledge the excellent technical support of the staff of TransViragen and the University of North Carolina Animal Models Core Facility.

